# Mechanism of Lutein to *meso*-Zeaxanthin Isomerization by RPE65 Catalysis

**DOI:** 10.64898/2025.12.10.693550

**Authors:** Alex Ballinger, Arunkumar Ranganathan, Maddison Bailey, Dylan Ramos, Annabel Lee, Binxing Li, Paul S. Bernstein, Martin P. Horvath

## Abstract

The macular pigments lutein (L), zeaxanthin (Z), and *meso*-zeaxanthin (MZ) protect the human retina from light and oxidative stress. While L and Z are abundant in the human diet, MZ is nearly absent. We previously demonstrated MZ is derived from L in chicken embryos precisely timed with expression of RPE65. Herein, we show that RPE65 from mouse, an animal that does not concentrate MZ in the eye, catalyzes L to MZ isomerization similarly as found for RPE65 from chicken and human, when expressed in cultured cells. Co-expression with xanthophyll-binding proteins had no impact on MZ yield. L and MZ both fit deep into the tunnel accessing the non-heme iron center, with strain evident for the *3’R*,*6’R* ε ring of L. Interestingly, a negatively charged Glu148, found along substrate tunnel, highly conserved among carotenoid cleavage dioxygenases, and which is critical for eye health, could be replaced by a neutral, isosteric residue (Gln) without impacting MZ yield. We propose that L to MZ isomerization proceeds by a neutral, radical transition state that differs from the carbocation encountered during retinoid isomerization. These findings extend our mechanistic understanding for macular carotenoid metabolism and should be considered when developing therapeutic interventions that act via RPE65.

## Introduction

The retinal pigment carotenoids (*3R, 3’R, 6’R*) lutein (L), (*3R, 3’R*) zeaxanthin (Z), and (*3R, 3’S meso*-) zeaxanthin (MZ) are distinctly concentrated in the foveal center of the macula in vertebrates, enhancing visual function and providing an additional layer of protection against blue light and reactive oxygen species ^[1]^. These damage-causing stressors are commonly associated with disorders of the retina, including age-related macular degeneration (AMD), glaucoma, Leber congenital amaurosis (LCA), and others ^[2]^. Over 750 carotenoids are found in nature, but notably, none of these molecules are physiologically synthesized *de novo* in vertebrates ^[1,3]^. Diets supplemented with L, Z, and MZ prevent visual loss from AMD and increase macular pigment spatial profiles ^[4,5]^. L and Z arrive at the retina through uptake in the gut, are then transported on HDL, and ultimately captured by xanthophyll-specific binding proteins expressed in eye tissues ^[6–8]^. While L and Z are abundantly found in fruits, vegetables, and other foods commonly found in the human diet, MZ is absent in the foods most people eat, present only in some marine animals. Sources of MZ include the skin and flesh of some fish, turtle fat, and shrimp carapaces ^[1,9,10]^, foods rarely eaten by humans, yet all healthy humans concentrate MZ in the retina.

A central question in retinal carotenoid biology is, where does the MZ found in the human retina originate? The *β* and *ε* ionone rings of L have the correct stereoconfiguration at C3′, matching the *β* rings of MZ, such that conversion requires only a π-bond migration from C4′–C5′ to C5′–C6′ without inversion of the tetrahedral center at C3′. This structural compatibility makes L a chemically plausible substrate for enzymatic isomerization to generate MZ ^[11,12]^, and has motivated investigations into the molecular basis of carotenoid metabolism within the retina. *In vivo* and *in vitro* studies support the idea that L is the precursor for MZ in primates and birds. When carotenoid-deficient monkeys were fed L, an increase in MZ was detected, but supplementation with Z had no impact on MZ ^[13]^. Japanese quail fed deuterium-labeled L generated stable-isotope labeled MZ in ocular tissues ^[14]^. These animal feeding experiments showed MZ is derived from L but did not identify the isomerase enzyme. Shyam et al. showed that the timing of RPE65-encoding messenger RNA expression matched MZ detection in developing chicken embryos, and that ocular MZ production could be abolished by an RPE65 inhibitor injected into the chicken egg ^[15]^. Additionally, heterologous expression of RPE65 orthologs from both chicken and human in HEK293T cells cultured in media supplemented with L produced MZ, whereas media supplementation with Z failed to generate MZ above background levels ^[15]^. These experiments strongly implicated RPE65 as the catalyst for L to MZ isomerization, and thereby identified a second activity for RPE65 ^[15]^, in addition to its primary function for retinoid isomerization (reviewed in ^[16]^; Figure 1).

**Figure 1.**
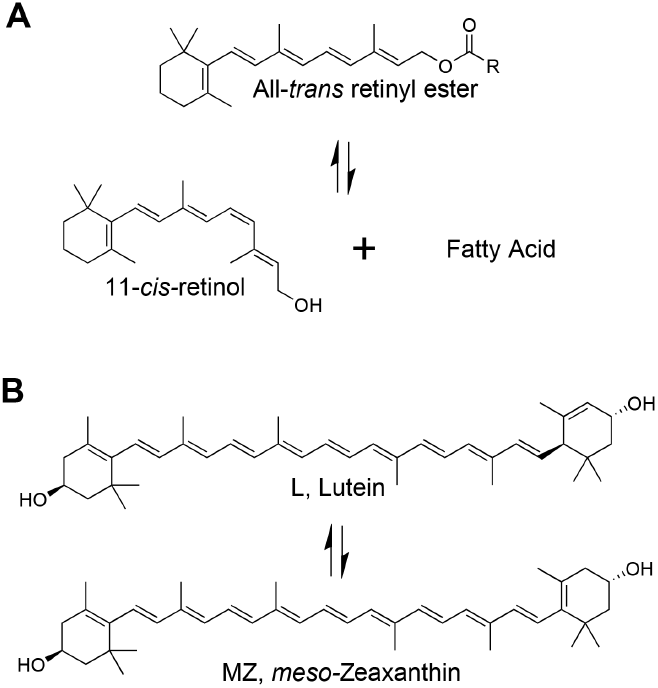
Reactions catalyzed by RPE65. The primary function of RPE65 is to catalyze retinoid isomerization, converting all-*trans* retinyl ester to 11-*cis* retinol and fatty acid by an atypical oxygen cleaving reaction (A). The secondary function of RPE65, and the focus of this report, is to catalyze L to MZ isomerization (B).

While the catalytic mechanism for retinoid isomerization in the visual cycle is well characterized ^[16,17]^, we know little about the catalytic mechanism for L to MZ isomerization. To address this gap, we further investigated L to MZ isomerization by RPE65 across human, chicken, and mouse orthologs, tested the impact of amino acid substitutions, and applied molecular docking to interrogate the enzyme-substrate and product complexes. Carotenoid metabolism mediated by RPE65 appears evolutionarily conserved, even for mouse, an animal that does not concentrate L, Z, or MZ in its retina ^[18]^. Differences in the yield of MZ observed for the different orthologs and the M450 variants found in inbred lines of mice are explained by differences in enzyme expression levels as detected by Western blot. Substitution of a negatively charged Glu at position 148 for its neutral isostere Gln (E148Q) had no significant impact on MZ yield and no discernable impact on protein levels, a surprising outcome given the absolute conservation of this glutamate, its importance for eye health, and the strong negative impact of the corresponding E150Q substitution for the apocarotenoid-cleaving oxygenase (ACO) from *Synechocystis* ^[19]^. We interpret these findings in terms of a proposed mechanism for RPE65-catalyzed L to MZ isomerization that proceeds by a neutral, radical transition state.

## Results and Discussion

### RPE65-catalyzed L to MZ isomerization is conserved across species

RPE65 requires membrane association for access to substrate and enzyme activity ^[20,21]^. To explore its mechanism, we and others employed transient transfection of cultured cells that normally do not express RPE65 ^[15,22,23]^. To test for L to MZ isomerization activity, cultured HEK293T cells were transfected with pCDNA3.1 plasmid DNA encoding RPE65 from three different species: chicken, human, and mouse. Special care was taken to ensure an identical CMV promoter context for each expression vector. The empty vector or the same vector with a gene encoding GFP in place of RPE65 provided negative controls for comparison. Cell culture media was replaced with media supplemented with 4 µM L 24 hours post-transfection. Cells were harvested 72 hours post-transfection, and carotenoids were extracted using a two-phase organic solvent separation, followed by drying under nitrogen and reconstitution in a 95% hexane / 5% isopropyl alcohol mobile phase (see Experimental Section). HPLC normal phase chromatography with a chiral column quantified MZ yield in relation to L as previously described ^[15]^. Comparable amounts of trace Z were detected in both experimental and negative control samples, suggesting a non-enzymatic or background origin.

Figure 2 summarizes cross-species comparison in terms of MZ yield, expressed as MZ/L. All RPE65-expressing cultures produced significantly noticeable MZ product relative to the GFP-expressing cultures (p < 3.0×10^−5^ for all comparisons), confirming that L to MZ isomerization is driven by RPE65 as previously reported ^[15]^. Chicken RPE65 (Gg WT) showed the highest normalized MZ levels, human RPE65 (Hs WT) generated somewhat lower MZ levels, and these orthologs were more robust relative to the two variants of mouse RPE65 (Mm L450 and Mm M450). The Mm L450 variant produced approximately twice as much MZ as the Mm M450 isoform (p = 1.1×10^-4^), an observation that mirrors differences in retinoid isomerohydrolase activity for inbred lines of mice expressing the L450 and M450 variants of RPE65, as reported previously by others ^[24,25]^.

**Figure 2.**
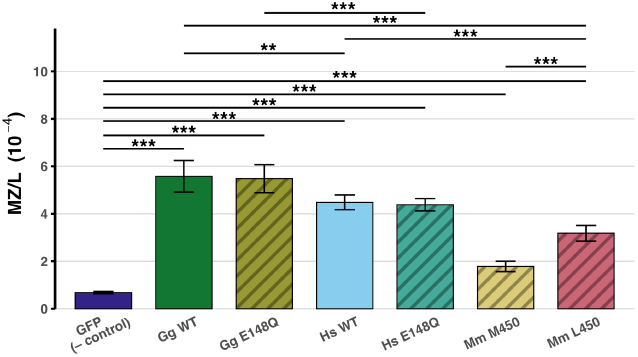
MZ product accumulation. MZ/L reports MZ product yield normalized by L substrate levels. Values are plotted as the mean of four biological replicates for cultured cells expressing different RPE65 orthologs and amino acid substitution variants grown in media supplemented with L. Species abbreviations: Gg, Gallus gallus (chicken); Hs, Homo sapiens (human), and Mm, Mus musculus (mouse). Error bars represent one sample standard deviation. Horizontal brackets indicate pairwise comparisons with annotations indicating statistically noticeable differences as determined by two-sided Student’s t-tests (* *p<0*.*05*; ** *p<0*.*005*; *** *p<0*.*0005*). All RPE65 orthologs and variants yielded statistically significant levels of MZ relative to the negative control cultures at the *p<0*.*0005* criteria. The Mm M450 and Mm L450 polymorphs yielded statistically noticeable different levels of MZ/L at the *p<0*.*0005* cutoff. Exact p-values relative to negative control are reported here: GgRPE65 *p = 1*.*7×10*^−*-6*^; GgRPE65 E148Q *p = 1*.*9×10*^−*7*^; HsRPE65 *p = 1*.*3×10*^−*7*^; HsRPE65 E148Q *p = 3*.*0×10*^−*8*^; MmRPE65 M450 *p = 2*.*9×10*^−*5*^; MmRPE65 L450 *p = 2*.*4×10*^−*6*^. For the comparison between MmRPE65 M450 and MmRPE65 L450, *p = 1*.*1×10*^−*4*^. For the comparison between GgRPE65 and HsRPE65, *p = 6*.*5×10*^−*3*^. For the comparison between GgRPE65 E148Q and HsRPE65 E148Q, *p = 3*.*4×10*^−*4*^. For the comparison between GgRPE65 and MmRPE65 L450, *p = 1*.*4×10*^−*4*^. For the comparison between HsRPE65 and MmRPE65 L450, *p = 3*.*9×10*^−*4*^.

To explore the source of differences in MZ production for the different orthologs, we evaluated protein expression levels by Western blot with an RPE65-specific antibody that recognizes the DALEED epitope conserved across all species tested (Figure 3). Total protein lysates were subjected to SDS-PAGE and transferred to nitrocellulose membranes, followed by immunodetection using a *β-actin-specific* antibody as a loading control, and with the DALEED antibody to detect RPE65 protein levels. Notably, probing with the RPE65-specific antibody detected a protein with mobility ∼65 kDa, intermediate to the 75-kDa and 50-kDa molecular weight markers, in all samples expressing RPE65, but failed to label the sample prepared from cultures expressing the GFP negative control.

**Figure 3.**
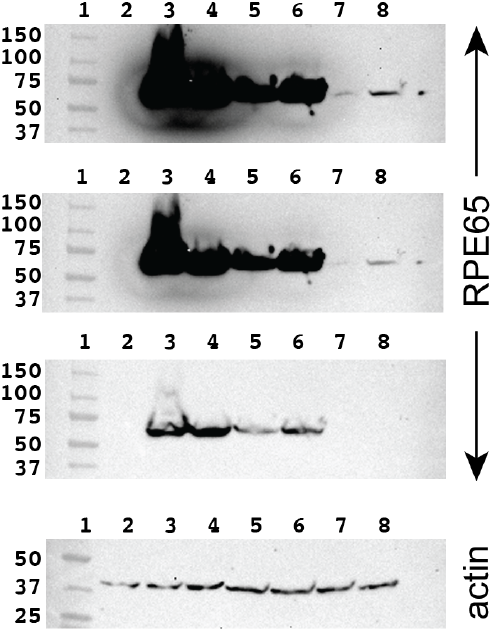
Protein expression levels. Cell lysates were denatured in SDS loading buffer and separated by PAGE prior to transfer to membranes and immunodetection. Top three images show three exposures for the same membrane probed with the anti-RPE65 antibody. The bottom image shows one exposure for the membrane probed with anti-*β*-actin. Samples were loaded left to right as follows: 1. molecular weight markers, 2. GFP negative control, 3. Gg RPE65 from chicken, 4. Gg RPE65 E148Q, 5. Hs RPE65 from human, 6. Hs RPE65 E148Q, 7. Mm RPE65 M450 from Mouse and 8. Mm RPE65 L450

This outcome confirms that our RPE65 expression system is working as intended and that HEK293T cells normally do not express the endogenous gene.

Strikingly, the trends in protein expression levels (Figure 3) match the trends for MZ/L levels (Figure 2). For example, expression of RPE65 from chicken is notably stronger compared to expression of RPE65 from human, and expression of RPE65 from mouse is very weak, a pattern that is also observed in the MZ/L values. The especially poor expression for the mouse RPE65 M450 polymorph explains why MZ yields are very low for cultures expressing this variant. These results are comparable to the previously described expression deficit in mice with the RPE65 M450 polymorph that explains lower visual pigment regeneration for M450/M450 mice relative to L450/L450 mice ^[26]^. Since the expression trends for mouse RPE65 appear conserved from animal to cell culture systems ^[24,26,27]^, and the only difference in our Mm RPE65-encoding genes is a single nucleotide polymorphism (SNP) at the first position of the (C/A)TG codon that encodes L450/M450, low expression of the RPE65 M450 enzyme may derive from differences in protein stability or differences in mRNA structure or stability consequent to the C/A SNP at codon 450 ^[28]^. From these observations, we suggest that RPE65 enzymes, even those from species that do not concentrate L, Z, or MZ in the retina, are intrinsically endowed with the secondary activity to isomerize L to MZ. The absence of MZ in *Bco2*^*–/–*^ mice, that otherwise accumulate L and Z ^[18]^, is therefore likely a consequence of enzyme insufficiency, and not the result of enzyme inactivity.

### Xanthophyll-binding proteins are not involved in the rate-limiting steps for L to MZ isomerization

To assess the potential influence of xanthophyll-interacting proteins on isomerization activity, we co-expressed human RPE65 with candidate proteins previously identified to transport or bind with L and MZ. In these co-expression experiments, pCDNA3.1 plasmid DNA encoding one of the proteins (all from human), GSTP1, SRB1, StARD3, or Aster-B, augmented the pCDNA3.1 plasmid DNA encoding Hs RPE65 for transfection in HEK293T cells. The pCDNA3.1 plasmid DNA encoding GFP served as a co-transfection control, meaning it was included together with the Hs RPE65-encoding DNA. Except for the addition of a second plasmid DNA, all procedures for transfection, timing of media supplementation with L, harvesting of cells, xanthophyll extraction, and HPLC analysis matched those for the single plasmid transfection experiments. Western blot analysis confirmed Hs RPE65 expression levels were not strongly impacted by including a second plasmid DNA. Table 1 compares MZ/L values for these co-expression tests with values measured for Hs RPE65 alone. Co-expression of RPE65 with other proteins diminished MZ yield somewhat, by about 10%, but this was true also for GFP, which is not expected to interact with RPE65 or xanthophylls. The modest reduction in MZ yield probably reflects competition for transcription and translation machinery. MZ yield under the condition of co-expression was remarkably consistent across the GFP (- control) and xanthophyll-binding proteins, suggesting that proteins known to interact with substrate L and product MZ neither assist nor inhibit the efficiency of RPE65-catalyzed isomerization of L to MZ. This outcome supports a model in which L accesses the active site of RPE65 via direct transfer from membrane-embedded reserves, consistent with the hydrophobic nature of L and the membrane-associated topology of RPE65. Of course, we cannot exclude the possibility of a protein-protein relay system to shuttle L and MZ to and from the RPE65 active site, but we are fairly confident that these steps, if in play, have no impact on the rate-limiting steps encountered during L to MZ isomerization.

**Table 1.** MZ yield with co-expression.

| Coexpressed protein | MZ/L ($10^{-4}$) * |
| --- | --- |
| none ‡ | 4.5 ($\pm 0.2$) |
| GFP (– control) | 3.9 ($\pm 0.2$) |
| GSTP1 | 3.9 ($\pm 0.1$) |
| SRB1 | 3.9 ($\pm 0.1$) |
| StARD3 | 4.0 ($\pm 0.3$) |
| Aster-B | 4.1 ($\pm 0.4$) |

\* The mean was calculated for three biological replicates.
Uncertainty is one sample standard deviation.
‡ Data without co-expression was calculated for four biological replicates.

### Molecular modeling of L and MZ complexed with RPE65

To visualize the substrate-enzyme and product-enzyme complexes associated with L to MZ isomerization, we docked these molecules into the active site of RPE65 with Autodock VINA ^[29,30]^. These experiments extend and significantly improve our previous molecular models derived by manual placement of L into the RPE65 substrate tunnel ^[15]^, because with Autodock VINA the search is more efficient, more exhaustive, and accommodates alternate residue rotamers. Xanthophyll ligands are abundantly found in the x-ray and cryo-EM structures of light-harvesting complexes and photosystems where lutein is packed closely with chlorophyll. These experimentally determined structures show a range of curvature for the polyene chain and characteristic dihedral angles at the junction with the ionone rings to minimize steric conflict for ring methyl groups. The non-zero dihedral angle is also observed for small molecule x-ray structures of L and MZ ^[31–33]^.

To sample the structural diversity expected for xanthophylls, docking was repeated for each molecule of an ensemble derived from experimentally determined molecular structures. The L and MZ ensembles correspond closely with the ligand with identifier LUT in the protein data bank ^[34,35]^. However, it was necessary to correct the *3S* configuration found in all LUT ligands to obtain the absolute configuration found for dietary L (*3R, 3’R, 6’R*) (see Supporting Information Experimental Section). As *meso*-zeaxanthin is not found in plants, MZ (*3R, 3’S*) molecules were constructed by grafting the *3S β* ring found in LUT molecules in place of the *ε* ring of L molecules. Ensemble members were treated as rigid ligands. Trials with flexible ligands or with ligands that allowed the *β* and *ε* rings to rotate were attempted but resulted in unrealistic structures with steric conflicts for the methyl groups attached to the ionone rings. The structure for bovine RPE65 in complex with difluoro-emixustat (PDB ID 7L0e) was selected as the experimentally determined receptor structure ^[36]^. Defined by x-ray diffraction to the 1.9-Å resolution limit, this structure is more accurate than AlphaFold-predicted models and yet appropriate for examining homologs since the 37 residues in contact with ligands are completely conserved across cow, chicken, human, and mouse.

Figure 4 shows representative poses for L and MZ characterized by nearly complete burial of the ligand in the emixustat-binding site of RPE65. One hydroxyl group is positioned near the mononuclear, non-heme iron center. The other ionone ring is located at the entrance to the deep tunnel. Emixustat is a transition state analog for the retinoid isomerohydrolase reaction catalyzed by RPE65. Comparing the structure of RPE65 complexed with difluoro-emixustat and palmitate (Figure 4A) with these docking outcomes (Figure 4B, Figure 4C and Figure 4D), we predict that L and MZ compete with the substrate for the retinoid isomerohydrolase reaction.

**Figure 4.**
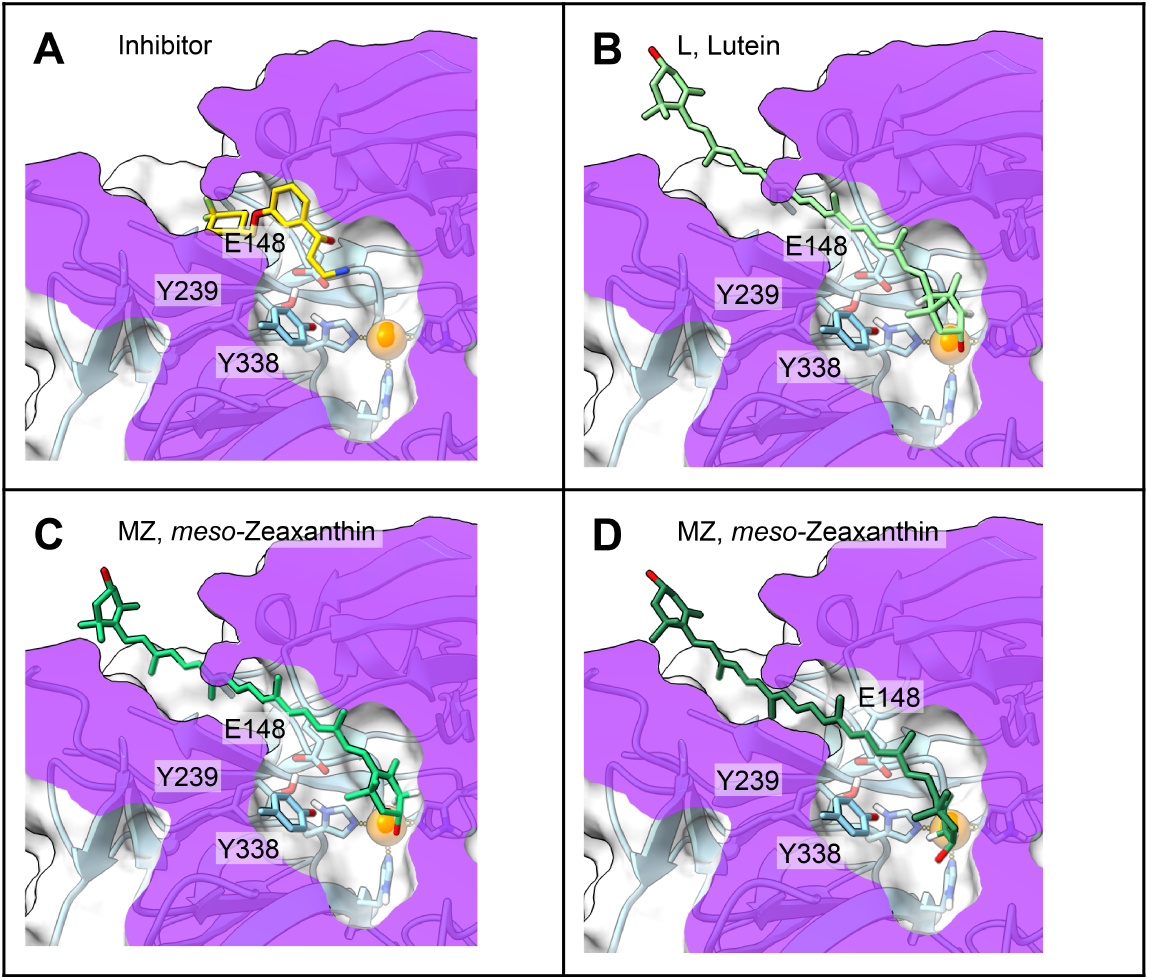
Molecular docking outcomes. The substrate access tunnel is revealed by clipping along a cross-sectional plane (purple). The active site of bovine RPE65 is occupied by the retinoid transition state analog difluoro-emixustat as shown in Panel A. Panel B shows a representative (*3R, 3’R, 6’R*) L docked in the substrate access tunnel. Panels C and D show two representative poses for (*3R, 3’S*) MZ docked with the OH group coordinating the iron center (Panel C) or disengaged (Panel D). The hydrogen atoms attached to C6’ in L and to C4’ in L and MZ are displayed for the L and MZ ligands. The H6’ hydrogen must migrate from C6’ in L to C4’ in MZ during isomerization.

Binding energies for the docked ligands revealed insights relevant for mechanistic understanding. Figure 5 plots the overall binding energy for different carotenoid ligands, limiting these to the top 10 winners with the *ε* ring of L or the corresponding *β* ring of MZ, Z, and *β-*carotene buried. The binding energies for these (*3R, 3’R, 6’R*) L “winners” were lower in magnitude compared to the binding energies measured for (*3R, 3’S*) MZ, (*3R, 3’R*) Z, or *β*-carotene, an outcome that indicates that the enzyme active site accommodates an *sp*^2^ center at atom C6’ more favorably than an *sp*^3^ center, and consequently exerts strain on the L substrate. Substrate strain was also evident from examination of orientation preferences. The (*3’R, 6’R*) *ε* ionone ring of dietary L was accommodated less frequently among the highest affinity outcomes compared with its (*3R*) *β* ionone ring, another manifestation that the chiral, *sp*^3^ hybridized C6’ center with tetrahedral geometry is more difficult to accommodate in the active site and therefore experiencing strain relative to an *sp*^2^ hybridized C6 center with trigonal planar geometry. For MZ, the *3R* and *3’S β* ionone rings were represented equally among the highest affinity docking outcomes, indicating that both R and S stereoconfigurations at the C3 and C3’ centers are accommodated without preference. We additionally tested docking with dietary Z and *β*-carotene and found no orientation preference, an observation that reassures us that orientation bias was not introduced consequent to construction of these ligands by computational methods.

**Figure 5.**
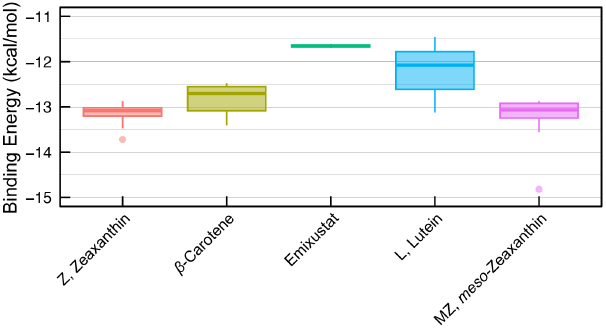
Binding energies for docked carotenoid ligands. The overall energy as evaluated by the “score only” option of VINA are plotted for the ten best outcomes with the *ε* ring of L or the corresponding *β* ring of MZ, Z, and *β*-carotene buried. More favorable binding outcomes have larger magnitude negative values. The overall winner among all docking outcomes is one MZ ligand with a binding energy of -14.8 kcal/mol. The most favorable outcome for an L ligand measured -13.3 kcal/mol. For comparison, the competitive inhibitor difluoro-emixustat measured -11.7 kcal/mol.

All of the top-10 (*3R, 3’R, 6’R*) L docking outcomes with the relevant *ε* ring buried adopted a highly similar pose as illustrated in Figure 4B. In this pose, the hydroxyl group was consistently coordinated with the mononuclear, non-heme iron center. Tyrosine residues could play an important role in a catalytic mechanism involving free radicals. Indeed, the hydrogen attached to C6’ in L, which must relocate to the C4’ carbon in MZ, was in close proximity to the OH group of Tyr 338 (Figure 4B). For the top-10 (*3R, 3’S*) MZ docking outcomes with the relevant *3’S β* ring buried, two poses were obtained (Figure 4C and Figure 4D). Eight of these outcomes were similar to the pose seen for L, with the hydroxyl group of MZ engaged with the iron (2.9-3.0 Å), and C6’ and H4’ about equidistant and relatively far from the OH group of Tyr338 (5.1 Å) (Figure 4C). Two of these MZ docking outcomes showed an alternate pose with the hydroxyl group disengaged from the iron center (Figure 4D). In this alternate pose, the hydroxyl group of Tyr 338 was closer to C6’ (3.4 Å) and closer to the H4’ hydrogen (3.9 Å) compared to distances observed for the first pose with the hydroxyl group coordinating the iron.

As evident from these docking outcomes, Glu148 comes into close van der Waals contact (3.4-3.8 Å) with the polyene chain of L and MZ and presumably is also in close contact with the transition state intermediates encountered during L to MZ isomerization. Glu148 is one of three absolutely conserved Glu residues that support three of the four His residues coordinating with the mononuclear, non-heme iron and which are thought to stabilize the positive charge on the iron center. Specifically, Glu148 hydrogen bonds with His241 and also with Tyr239 (Figure 4). It is assigned a formal negative charge by the *addcharge* module of ChimeraX, thus imparting a negative electrostatic potential to this region of the substrate tunnel.

### The transition state for RPE65-catalyzed L to MZ isomerization is uncharged

The close proximity of Glu148 to the substrate tunnel of RPE65 suggested a means to probe the nature of the transition state encountered during isomerization of L to MZ. We changed the codon for residue 148 in the genes encoding the RPE65 orthologs from chicken and human to replace Glu148 with Gln (E148Q) and thereby neutralized the negative charge at this position, while retaining hydrogen-bonding capacity and shape. The MZ yield obtained with cultures expressing each of these E148Q variant enzymes measured well above background values, indicating catalysis of L to MZ isomerization (Figure 2). Indeed, MZ yield was unperturbed, within experimental uncertainty, by the E148Q replacement for both the chicken and human enzymes. For example, the wild-type Hs RPE65 from human yielded MZ/L of 4.4 (± 0.2) × 10^-4^, and the E148Q variant yielded MZ/L 4.3 (± 0.2) × 10^−4^. Likewise, the wild-type Gg RPE65 from chicken yielded MZ/L of 5.6 (± 0.5) × 10^-4^, and its E148Q variant yielded MZ/L of 5.8 (± 0.3) × 10^-4^. Western blot analysis probing with the anti-RPE65 antibody confirmed comparable protein abundance for each wild-type and E148Q variant (Figure 3). These results show that Glu148, despite being absolutely conserved among RPE65 orthologs and carotenoid cleavage oxygenases, can be replaced with its neutral, isosteric carboxamide analog without altering protein stability or impacting L to MZ isomerization.

This is a surprising outcome given that replacement of Glu148 causes retinal pathology, leading to Leber congenital amaurosis in humans ^[37]^. The corresponding E150Q replacement for the apocarotenoid cleavage oxygenase (ACO) from *Synechocystis* strongly impaired catalysis with an 80-fold reduction in *k*_*cat*_*/K*_*M*_, further reinforcing the mechanistic importance of this residue ^[19]^. While the negative charge of this Glu residue provides critical functional properties for RPE65 in visual pigment regeneration, and for carotenoid cleavage oxygenases, consistent with a carbocation transition state for these reactions ^[16,17,38–41]^, the negative charge is dispensable for the second function of RPE65. For a charged transition state, the rate of isomerization would be expected to dramatically increase or decrease with the E148Q substitution because electrostatic interactions either destabilize a negative transition state or stabilize a positive transition state. Our finding that MZ product yield is unchanged upon E148Q substitution for two different RPE65 orthologs strongly argues against significant charge on the transition state for L to MZ isomerization, as catalyzed by RPE65.

The clinical relevance of Glu148 is supported by genetic and mechanistic studies identifying this residue as a sensitive determinant of RPE65 function. Early genetic analyses demonstrated that pathogenic substitutions within the mononuclear iron’s microenvironment cause severe early-onset retinal dystrophy ^[42]^. Subsequently identified, E148D is among disease-associated variants in LCA patient cohorts, highlighting that even conservative substitution at this position can produce clinically significant pathology ^[37]^. These genetic patterns align with structural insights showing that Glu148 forms part of the second-sphere network integral to tune metal center character and stabilize reaction intermediates (reviewed in ^[16]^). Mutational studies of this four-His/three-Glu architecture reinforce this view, showing that replacement of conserved glutamate or histidine residues abolishes isomerohydrolase activity ^[43]^. By extension, Glu148 is likely essential for the canonical retinoid isomerization function of RPE65, yet the acidic character of Glu148 is unnecessary for catalysis of L to MZ isomerization. The E148Q replacement thus separates the two functions of RPE65, and leads to the idea that the two chemistries proceed through distinct transition states.

### A radical mechanism for RPE65-catalyzed L to MZ isomerization

Our investigation of the E148Q substitution variants of RPE65 points to an uncharged transition state for L to MZ isomerization, with the further inference that free radicals are involved. Here, we integrate these ideas with insights from molecular models to propose a plausible radical mechanism for the conversion of L to MZ. RPE65 is endowed with structural features that are well-suited for creating and trapping a free radical, such as the iron metal center, capable of multiple oxidation states, and several nearby tyrosine residues, which are well-known to play a central role in storing radical intermediates. Indeed, the transition state for retinoid isomerization and carotenoid cleavage is discussed in terms of a carbocation radical intermediate ^[38–40]^, setting a precedent for radical mechanisms catalyzed by this family of enzymes.

Figure 6 illustrates our proposal for a noncanonical isomerization mechanism with neutral, radical transition state intermediates. Initially, hydrogen (H•) moves from C6’ of the substrate L to a radical tyrosyl at position 338, to create the neutral xanthophyll radical intermediates pictured in panel 2 and panel 3 of the mechanism.

**Figure 6.**
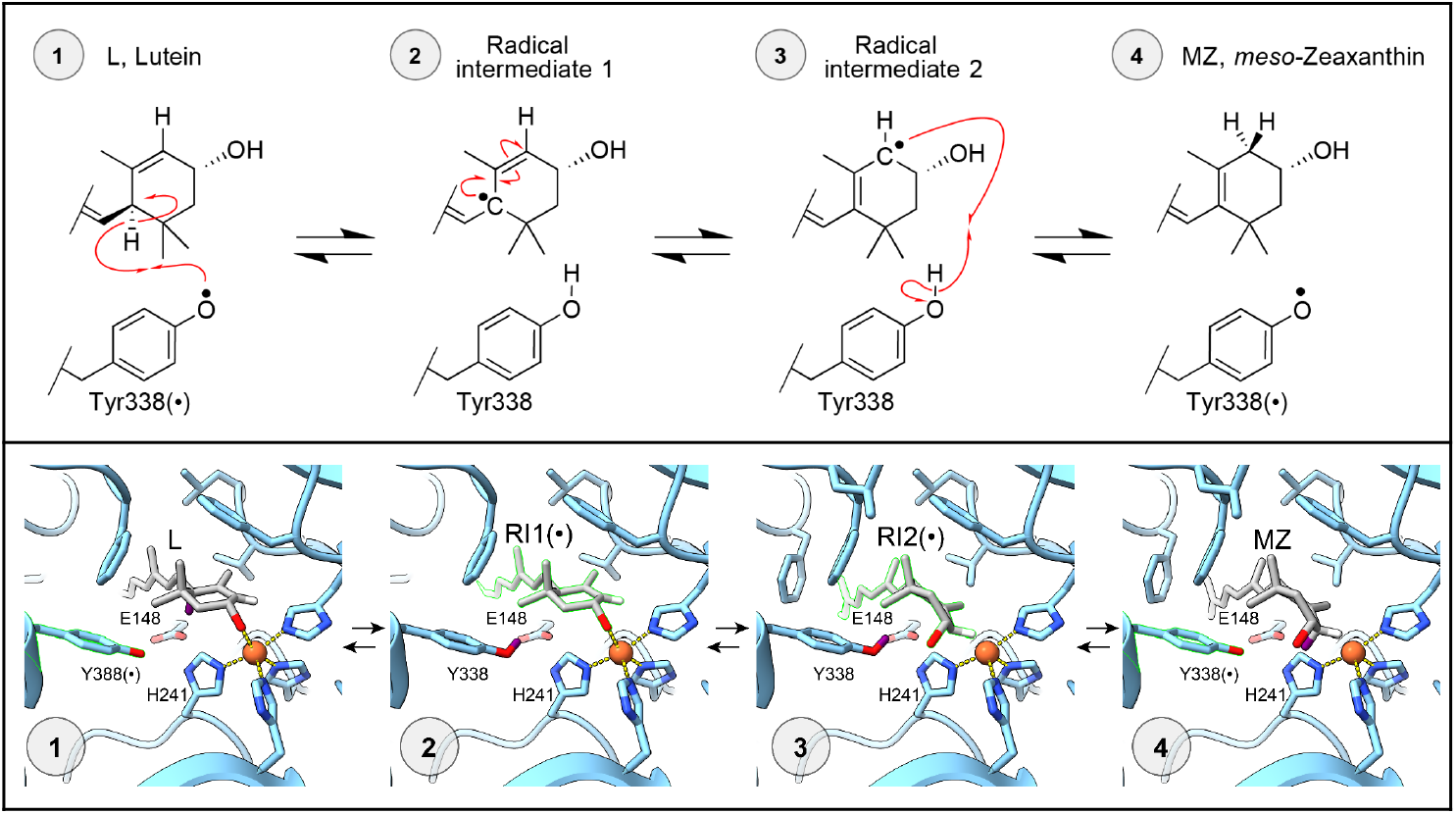
Proposed mechanism for L to MZ isomerization catalyzed by RPE65. A Tyr338(•) radical (panel 1) is generated by proton-coupled electron transfer, possibly at a large distance from within the RPE65 dimer. H• transfer from C6’ of L to Tyr338 creates the uncharged free radical intermediates with the radical on C6’• (panel 2) in equilibrium with the radical on C4’• (panel 3). H• transfer from Tyr338 to C4’• creates MZ (panel 4).

Subsequently, the H• moves from Tyr338 to C4’ to effect selective double bond migration and generate the product MZ. Molecular oxygen is not involved directly in this mechanism as the hydroxyl group of the ionone ring is coordinated with the iron center, at least initially. However, the radical tyrosyl may ultimately be derived from reactive oxygen species encountered prior to entrance of L into the substrate tunnel or at the second iron of the RPE65 dimer.

Our proposed mechanism builds upon radical chemistry in non-heme iron enzymes such as lipoxygenases, which abstract bis-allylic hydrogens through Fe(III)-superoxo intermediates and proceed via allylic radical rearrangements ^[44,45]^. Similarly, Fe(II)/Fe(IV)-dependent halogenases like SyrB2 catalyze substrate-centered radical rebound on lipid-like scaffolds ^[46]^, supporting the feasibility of such chemistry in hydrophobic environments. This is seen in the similar carotenoid oxygenase NinaB, in which the cleavage of the carotenoid substrate is activated by a superoxide intermediate generated at the enzymatic iron cluster, which can displace a free radical through a conjugated network ^[47]^. The original source of the free radical and its final quenching may involve similarities to proton-coupled electron transfer (PCET), as found for mechanisms proposed in photosystem II, ribonucleotide reductase, and flavin monooxygenases, where concerted proton and electron transfer is mediated through hydrogen-bonding networks involving conserved Tyr, Trp and His residues ^[48–50]^. In addition to transiently stabilizing the radical, geometric restriction necessary for selective isomerization may also be accomplished by these aromatic residues ^[51]^. The homodimeric architecture of human RPE65 ^[52]^ suggests that inter-subunit PCET may link the two active sites, as is the case for class I ribonucleotide reductase, where long-range radical transfer occurs across *α*2–*β*2 subunits ^[50,51]^.

The proposed mechanism bypasses the need for full carbocationic or carbanionic intermediates, offering a chemically and physiologically compatible pathway for MZ production within the oxygen-rich environment of the retinal pigment epithelium ^[53]^. To further explore this radical mechanism, the source and location for short-lived intermediates should be mapped by high-resolution crystallography paired with electron paramagnetic resonance of crystallized samples, as shown for other enzymes ^[54]^. Site-directed mutagenesis of Tyr, Trp, and His residues would help define the PCET network. These strategies must be adapted for RPE65, an enzyme that is notoriously sensitive, not easily purified, acts on substrates that are insoluble in water, and strictly depends on a native lipid environment ^[21,55]^. Characterizing a radical-mediated pathway would provide a mechanistic understanding for reduced MZ associated with variants of RPE65 in patients with AMD and other retinal degenerations.

### Clinical significance

Mutations in RPE65 associated with inherited retinal diseases can provide mechanistic insights. As the third member of the mammalian carotenoid cleavage dioxygenase superfamily that is critical for normal vision, its enzymatic mechanism of action as a retinoid isomerohydrolase has been studied extensively on a molecular level and correlated with known human mutations ^[16]^. Because RPE65 has two activities, we may expect three classes of mutations: a class similar to L450M that reduces enzyme expression or stability and therefore impacts both retinoid isomerohydrolase function and L to MZ isomerization; a class similar to Glu148 replacement that disables retinoid isomerohydrolase activity but leaves L to MZ isomerization intact; and possibly a third class that impairs L to MZ isomerization but supports normal visual pigment regeneration. From our investigation of the impact of Glu148 replacement on L to MZ isomerization, we expect normal MZ levels for LCA patients with poor visual pigment regeneration capacity consequent to E148D replacement. Unfortunately, distinguishing the identities of the three macular pigment carotenoids cannot be done noninvasively and requires postmortem eye donation, so direct proof of this hypothesis is unavailable.

The highest concentration of the macular carotenoids is at the foveal center, and it is well established that the fovea is particularly enriched in MZ and Z relative to L ^[56]^, yet Z is 5-10 times less common in the diet, and MZ is essentially absent ^[1]^. Biochemical mapping studies established that M and MZ are concentrated at the foveal center, forming a gradient that decreases toward the periphery, while L shows the opposite gradient, increasing toward the periphery ^[56]^. High-resolution resonance Raman imaging shows that the foveal pigment is up to 90% Z + MZ, while L is more evenly distributed across the retina at a much lower concentration ^[57,58]^. Reciprocal gradients of the zeaxanthins *versus* L fit with the idea that MZ is locally generated from retinal L to increase the carotenoid levels in the fovea beyond levels achievable by dietary zeaxanthin alone ^[58]^. Biochemical analyses further show that MZ has superior capacity for singlet oxygen quenching compared to L and Z, and that mixtures containing all three carotenoids quench more efficiently than any individual component ^[59]^. Taken together, data support the view that central macular pigment relies critically on MZ and that disruptions of L to MZ isomerization could have detrimental effects on visual function and physiology.

## Conclusion

First described as an activity fractionated from frog retina/pigment epithelium homogenates that transformed all-*trans* retinol into 11-*cis* retinol ^[60]^, the mechanism of visual pigment regeneration, and the central role played by RPE65 in retinoid isomerization is now well understood ^[16,17,61]^. More recently, RPE65 was identified as the enzyme catalyzing L isomerization to explain how the non-dietary xanthophyll MZ is derived in retinas of primates and birds ^[15]^. Here, we expand our understanding of this secondary function of RPE65. Our experiments extend the range of RPE65 orthologs from vertebrate species previously tested, confirming that RPE65 possesses a second enzymatic function catalyzing the isomerization of L to MZ, even for RPE65 in animals adapted to low light. These results align with improved molecular models, constructed with realistic structures for xanthophylls that correct a misassigned stereoconfiguration prevalent in databases, showing substrate L and product MZ are both accommodated in the retinoid-binding tunnel of RPE65, where residues are completely conserved.

Species-specific differences in RPE65 are located on the surface of the protein, far away from the iron center, its ligand sphere, and this substrate access tunnel. The transition state for L to MZ isomerization is different from that of retinoid isomerization, as inferred from full retention of MZ yield for E148Q substitution variants of RPE65 from both chicken and human. As far as we know, the E148Q substitution has not previously been attempted for RPE65. The corresponding E150Q replacement in ACO from *Synechocystis* strongly impaired carotenoid cleavage ^[19]^, and replacement of Glu148 with Asp is associated with Leber congenital amaurosis consequent to failed visual pigment regeneration in human patients ^[37]^. While highly destabilizing for a carbocation transition state, the removal of negative electrostatic potential by E148Q replacement would have little impact for an uncharged, radical transition state. This reasoning leads to a proposed radical mechanism for RPE65-catalyzed L to MZ isomerization that proceeds with H• migration first from C6’ of L to a tyrosyl radical to create the uncharged radical xanthophyll transition state intermediate, followed by transfer of H• to C4’ to generate MZ. Our findings expand the biochemical function of RPE65, and reveal that mutations predicted to disable retinoid isomerization leave the MZ to L isomerization function of RPE65 intact. Therapeutic interventions that competitively inhibit RPE65 should consider the side-effect of reduced MZ production predicted by our work.

### Experimental Section

See Supporting Information for the Experimental Section.

## Data and Reagent Availability

Plasmid DNAs have been archived with AddGene. The binding energies and other data describing carotenoid docking outcomes are archived with DataCite. Scripts, programs, and molecular structures for accomplishing the docking experiments are archived with GitHub.

## Acknowledgements

This work was supported by National Institutes of Health Grants EY11600 and EY14800 to P.S.B., and an unrestricted departmental grant from Research to Prevent Blindness. We thank Dr. Jian-Xing Ma for generously providing the plasmid DNAs encoding chicken RPE65 and human RPE65. Molecular graphics and analyses performed with UCSF ChimeraX, developed by the Resource for Biocomputing, Visualization, and Informatics at the University of California, San Francisco, with support from National Institutes of Health R01-GM129325 and the Office of Cyber Infrastructure and Computational Biology, National Institute of Allergy and Infectious Diseases. The support and resources from the Center for High Performance Computing at the University of Utah are gratefully acknowledged.

## Supporting Information

The authors have cited additional references within the Supporting Information ^[62–65]^.

