## Supporting Information for "Mechanism of Lutein to *meso*-Zeaxanthin Isomerization by RPE65 Catalysis"

**Figure S1.** Uncropped Western blot images

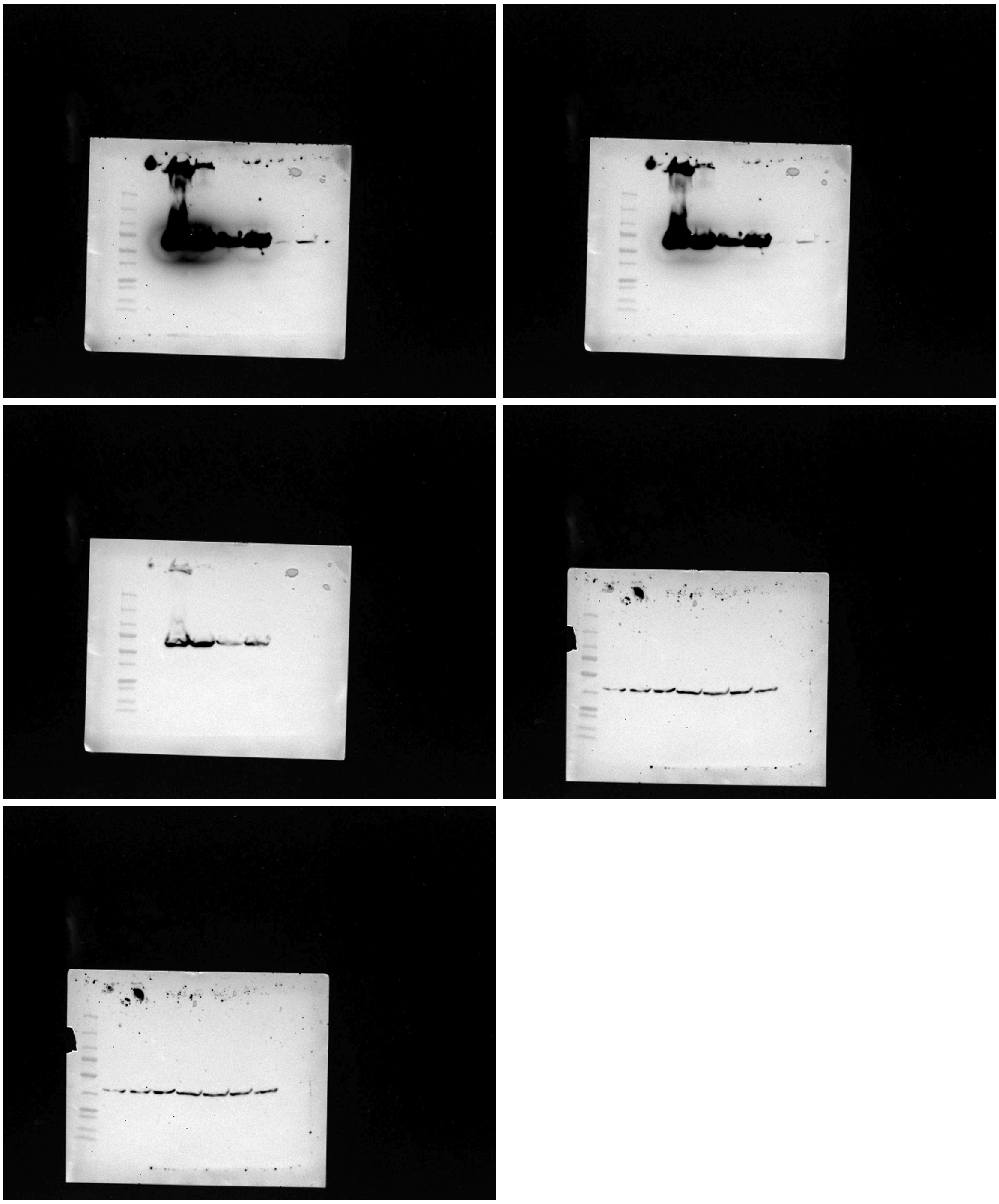

**Gene identifiers for RPE65**

**Table S2**. Gene accession identifiers for cloning

| Species (variant) | GenBank Accession ID | Modifications | Modification Type |
|---|---|---|---|
| Gg RPE65 | AB017594.1 | C318>T, G606>A, T963>C, C1156>T, C1161>T | Silent SNPs |
| Gg RPE65 (E148Q) | AB017594.1 | *ibid* G441>A, G442>C | *ibid* E148Q Mutation |
| Hs RPE65 | U18991.1 | T465>C | Silent SNP |
| Hs RPE65 (E148Q) | U18991.1 | *ibid* G442>C | *ibid* E148Q Mutation |
| Mm RPE65 (M450) | AF410461 | None | None |
| Mm RPE65 (L450) | AF410461 | A1348>C | M450L Mutation |
| GFP | PV533970.1 | C681>T, C684>T | Silent SNP |
| GSTP1 | X06547.1 | G632>A, A633>G | Silent SNP |
| SRB1 | KJ901323.1 | None | None |
| StARD3 | D38255.1 | None | None |
| Aster-B | NM_020716.4 | Numerous* | Optimized for bacterial expression |

* See sequence provided below

**Aster-B Nucleotide Sequence codon optimized for bacterial expression**
atggtatcggtttctcttctcctcactctcgcttgtataatcccgagacctaataccgttttgaaggttgatcaggctatctttcatttgatccaggagcattacggagctct
ttattatctctcgccacttctgcagctctgtatcatgatatttctgctgactctcaagcagctgcgcccattcggtttgagactgtggaagtctctcctgcagcctcagcc
cttgccaggcggtgagcgtttgtgttgtatactcgagcatccaaagtttatagaacagcatcatattaagaatgaccaacaggaccagggaaaagcatatcaca
caggaaataaccaaaaggagttttgaaacactttgcagatggaagccatttggtgtatcttcaggaatgtggcgggtttgggtggagccagccacatgctttattct
atggcctacgtcctcatccgtaggcgtggtaaccggagacatgacttcctccaagtgagggactctaaggtgagcgtgcggtcttttgcggcggcgaaccgtag
tggtttttgatgccttttctttaggagactgcctatgcatttcagcaagataagtggattctgtcttagccagttcgctttcgaggtggcggaagtagtcttccaggccgg
accaaaaattcttctcgataaaagtcttcaccagtccccaaggttgttttcggtaccgcaattccgtgctaaccctaagtcgggatttattcctcgcaactcgcgtca
gggtatagcgattgatggtatagaagtaatcgtgataaggcacatcatgggtaagcacctcggcgtcaatcacgtagcattctgactcctgactggctttgtacat
cgtctgagtctctctcacggttgcagtcttgggagccaaaggatttgtcaaggttatggtgtacaatatgacgcgggattggttaccgttctcttctttttccatgggtg
gaaaattatgtcgctgaatctgcgttgttccataaagtccctctggaacggagagttcgtgaaaaggagatcataaagtttgtcaacgctaaaattgaaaacttcat
tcacgtattgtcgaccagagagatcttcgtaaaaggcctggacctcgccctcatcgtgggtatcactagaatcgctaagctcggttggtatatcttcgttatcattaa
aatcgagtgagggactgtttacgggagctatcatttcgatcttttcgcccatgatgttatcgatagcgagttctttctccagggagccgtcaccctcaagagcctcttct
tcaagaggcagcccatcaaaggaaactggagcttcgctactgccggtgctagtcagggtactattcgtgattgatttttaggcagttgaggtgacgcatcgggttt
tgtctcgatggaagacttggaagaagaatcatttacctcattttcctccacaggaatctcctcacaataacccattgtattaaagtcgtcatcgggcggaacgtagt
cttcgtcatcggaagtgagaccgagctcattaccatagcactgatgaacaaaatgccagagttcctttgggcacaagggtttctccaggagagcgttttgccaaa
ggcgaaacatcatcatgtaggtcctgtcgcgggcaccgaaagaagtgaagaaatgtttctcactgtcagtacaaacctgtattgcattgggaataagcctagcg

gtcttctctttcgtcatactgcaaatatccttcaatcggacagtcaggagagtctcccacctaaagatgtttgaatagaagcaaatccaattctcggaaagatacag
ccgcccctgcagcaggatgtctcgttggagtgcgcatgagtagtctacgatcaaccgctctgtatctgggagctgcttgaacaattttctaaaatcttcgttcctttgttt
ataggtcgggctcaagacgttgtaccaggattgtgacttcttgctattcttcccaccgcttctaccggattggctggagcatgagcgttgcacccccttgctctggagtg
cttggacttttatcagaggagtggtctgaacccttctcaaccattgtgtctccatcctgattttgtggtgtcggggaccggctgcgtttcctcagaattgggctacacgc
cggtgtacttctgttactatttgaggccgtgcaactaagtttgaatcccttcat

### Experimental Section

#### Molecular cloning

Expression constructs encoding human, chicken, and mouse RPE65 variants and xanthophyll interacting proteins were generated by standard molecular biology methods, starting from a chicken RPE65-encoding pCDNA3.1 vector from the Jian-Xing Ma group, with a cytomegalovirus (CMV) promoter for amplified expression in mammalian cells.

The pCDNA3.1 vector was linearized by digestion with *NotI* and *HindIII* (New England Biolabs), followed by dephosphorylation with Antarctic Phosphatase (New England Biolabs) to minimize vector self-ligation, and the 5361-bp DNA was purified by agarose gel electrophoresis with the GeneJet Gel Extraction Kit (Thermo Fisher Scientific) according to the manufacturer's instructions. Coding sequences for RPE65 variants from human, chicken, and mouse, as well as xanthophyll interacting proteins, including wildtype Hs RPE65 (accession ID U18991.1), Hs RPE65 E148Q (U18991.1 G442>C), Gg RPE65 (AB017594.1), Gg RPE65 E148Q (AB017594.1 G441>A+G442>C), Mm RPE65 Met450 (AF410461.1), Mm RPE65 Leu450 (AF410461.1 A1348>C), GFP (PV533970.1), GSTP1 (X06547.1, G632>A+A633>G), SRB1 (KJ901323.1), StARD3 (D38255.1), and Aster-B (NM_020716.4) were synthesized as PCR products by amplifying template genes already available or templates provided by synthetic DNA Gene Blocks (Integrated DNA Technologies) or combinations of these templates. PCR products included *NotI* and *HindIII* restriction endonuclease recognition sequences nearby the 5' and 3' termini. PCR products encoding mouse RPE65 consistently yielded clones with missense mutations. This challenge was remedied by replacing the template with a HiFi Gene Block (Integrated DNA Technologies) and screening several clones. In those cases for which the target gene contained internal *HindIII* or *NotI* recognition sequences, these were removed by synonymous single nucleotide substitution. These synonymous nucleotide substitutions and the nucleotide substitutions needed to replace Glu148 with Gln (E148Q) or replace M450 with Leu (M450L) were introduced by PCR with overlapping primers carrying the altered nucleotide sequence. The DNA encoding the target gene was purified by agarose gel electrophoresis, digested with *NotI* and *HindIII,* and repurified with the GeneJet Gel Extraction kit to remove the restriction enzymes to generate the insert DNA for ligation reactions. Insert DNA and linearized pCDNA3.1 DNA were ligated using Quick Ligase and Quick Ligase Buffer (New England Biolabs) following the manufacturer's instructions.

Ligation reactions (2 µL) were mixed with competent *E. coli* DH5α cells (50 µL) in a thin-walled PCR tube, incubated on ice for 3 min, and heated at 42°C for 45 s. The heat-shocked cells were rescued by incubation in SOB media at 37 °C with shaking for 45 min, and transformants were selected on LB agar plates supplemented with 100 µg/mL ampicillin. Colonies were purified and subsequently cultured in LB broth for plasmid preparation using the Plasmid Miniprep System PureYield (Promega). Plasmids were subjected to whole-plasmid sequencing performed by Plasmidsaurus using Oxford Nanopore Technology with custom analysis and annotation. Sequencing reads were aligned to reference plasmid sequences using Clustal Omega to confirm authenticity. Plasmid DNAs with RPE65-encoding sequences and correct vector architecture were selected for HEK293T transfection.

#### HEK293T jetPRIME transfection

To evaluate the enzymatic activity of RPE65 in mediating the isomerization of lutein to *meso*-zeaxanthin, we employed the HEK293T mammalian cell line, an immortalized human embryonic kidney derivative that lacks endogenous expression of RPE65. HEK293T cells were cultured in high-glucose DMEM (ATCC 30-2002) supplemented with 10% (v/v) fetal bovine serum (ATCC 30-2020), 1% (v/v) streptomycin/penicillin (Gibco™ 14040133). For transfection experiments, approximately 1.5 x $10^6$ HEK293T cells were seeded into a T75 flask containing 10 mL DMEM culture medium and transfected after achieving 70-80% confluence, typically in 24-36 h. All transfections were carried out using a jetPRIME Transfection kit following jetPRIME's *Short Protocol - DNA Transfection* guide. For single plasmid experiments, plasmid DNA (10 µg) was mixed with jetPRIME buffer (500 µL), vortexed for 10 s, and centrifuged for 30 s. JetPRIME reagent (20 µL) was added to the solution, vortexed for 10 s, and centrifuged for 30 s. The solution was added dropwise evenly to a 10 mL flask of seeded cells. For co-expression transfection experiments, the volumes were doubled to account for the additional DNA (10 µg from the human RPE65-encoding plasmid and 10 µg of a second plasmid). The combined DNA was mixed with jetPRIME buffer (1 mL), vortexed for 10 s, and centrifuged for 30 s before jetPRIME reagent (40 µL) was added. Transfected cells were incubated in 5% $CO_2$ at 37° C with no light to protect carotenoids from ambient light.

#### Supplementation with substrate L

Twenty-four hours post-transfection, the media was removed and replaced with 10 mL fresh DMEM media, and L, previously suspended at a concentration of 100 µM in serum-free DMEM with 0.01% concentration TWEEN 40 (Sigma Aldrich: P1504), was added evenly across cells in the flask to deliver a final 4 µM concentration of L. L was verified by HPLC to be free of MZ. Cells in L-supplemented media were incubated for an additional 72 h in 5% $CO_2$ at 37° C in the dark.

**Extraction of carotenoids**

Cells were harvested 96 h post-transfection. To accomplish this, the DMEM media was removed, and cells were detached using 3 mL TrypLE (Gibco TrypLE™ Select Enzyme (1X), no phenol red) and collected by low-speed centrifugation at 300 × *g* for 5 min. The supernatant was removed, and cells were washed by vortexing with 3 mL PBS buffer (Gibco™ DPBS, calcium, magnesium). Cells were collected by low-speed centrifugation at 300 x *g* for 5 min, and the supernatant was removed. The washing process was repeated a total of three times. Carotenoids were extracted from the washed, transfected HEK293T cells via a biphasic organic/non-organic separation protocol optimized for hydrophobic pigment isolation [62]. In a dark room at ambient temperature, 1 mL of tetrahydrofuran with 0.01% (w/v) butylated hydroxytoluene (THF•BHT), a solvent known to efficiently solubilize carotenoids while minimizing degradation, was added to cells and the mixture was homogenized by sonication for 10 min, vortexed for 30 seconds, and membranes and insoluble materials were pelleted by centrifugation at 13,000 × *g* for 5 min. The THF•BHT-soluble phase, with a bright yellow color indicative of carotenoids, was collected, and the pellet was set aside. The collected liquid was dried using a gentle stream of nitrogen gas to prevent oxidative degradation. The cell pellet was re-extracted by resuspension in 1 mL THF•BHT. The soluble extract was combined with the previously extracted sample and evaporated under nitrogen. Re-extraction was repeated until no visible yellow color was apparent in the solvent, typically 3 or 4 cycles. The dried material was subject to a secondary liquid-liquid extraction by adding sequentially 200 µL ultrapure water, 400 µL methanol, and 600 µL hexane. The resulting mixture was sonicated for 10 min, vortexed for 30 seconds, and centrifuged at 13,000 x *g* for 5 min to achieve clear separation of layers. The upper yellow hexane layer was carefully collected and dried under nitrogen. Extraction with hexane (600 µL) was repeated and combined with the previously extracted sample until no visible yellow color was apparent in the solvent, typically 3 or 4 cycles.

**Separation and quantification by normal phase HPLC**

Dried extracts were reconstituted in a mobile phase consisting of hexane:isopropanol (95:5, v/v) and injected for isocratic separation and xanthophyll quantification. Samples were analyzed on an Agilent 1260 Infinity II HPLC system equipped with a photodiode array detector. Carotenoids were resolved using a normal-phase chiral column (Chiralpak AD, 250 × 4.6 mm, 10 µm, Chiral Technologies, West Chester, PA) fitted with a 50 x 4.6 mm column guard. The mobile phase was hexane:isopropanol (95:5, v/v), set to a flow rate of 0.7 mL/min, with a total run time of 60 min per sample. Peaks corresponding to L and MZ were identified by spectral signatures, quantified by peak area and normalized as the MZ/L ratio, as previously described [15].

**Western blot**

Cells and tissues were homogenized on ice in RIPA buffer (25 mM NaCl, 0.5 mM EDTA, 25 mM Tris·HCl, pH 7.2, and 0.1% v/v Tween-20) supplemented with protease inhibitors. Lysates were cleared by centrifugation, and protein concentrations were quantified using the BCA assay (Thermo Fisher Scientific). Equal amounts of protein (20 µg) were separated by 9% SDS–PAGE and transferred to 0.45-µm nitrocellulose membranes (Bio-Rad Trans-Blot SD, 25 V, 1 hour). Membranes were washed in Tris-buffered saline (20 mM Tris·HCl, 500 mM NaCl, pH 7.5) supplemented with 0.01% Tween-20 and blocked for 1 h in Odyssey Blocking Buffer (LICORbio) supplemented with 0.01% Tween-20.

Primary antibody incubation was performed overnight at 4 °C using a 1:1,000 dilution of the monoclonal anti-RPE65 antibody DALEED (Thermo Fisher Scientific Catalog # MA1-16578; Clone ID 401.8B11.3D9). A mouse monoclonal anti-*β*-actin antibody (8H10D10; Cell Signaling Technology, 1:5,000) served as a loading control. Following incubation with HRP-conjugated secondary antibodies (1:10,000, 1 hour at room temperature), signals were detected using the Invitrogen™ iBright™ FL1500 Imaging System with several exposures. The pre-stained molecular weight markers were obtained from BioRad (Catalog #161-0374).

**Correction of stereoconfiguration for dietary L**

Many examples of molecules closely corresponding with dietary L but with an inverted C3 center (*3S, 3'R, 6'R*) are found in the Protein Data Bank [34,35] with the ligand identifier LUT. An ensemble of 93 LUT ligands determined to 2.8 Å resolution limit or better were harvested from the x-ray crystal structures and cryo-EM structures of light-harvesting complexes and photosystems from plants and cyanobacteria. As these molecules are found in the dietary source of L, we believe that the stereoconfiguration for LUT was misassigned in the original ligand description and then propagated throughout structures deposited with the Protein Data Bank. Misassignment of stereoconfiguration is made more likely since the *R* and *S* designations are determined with xanthophyll-specific priority assignments (Figure S3).

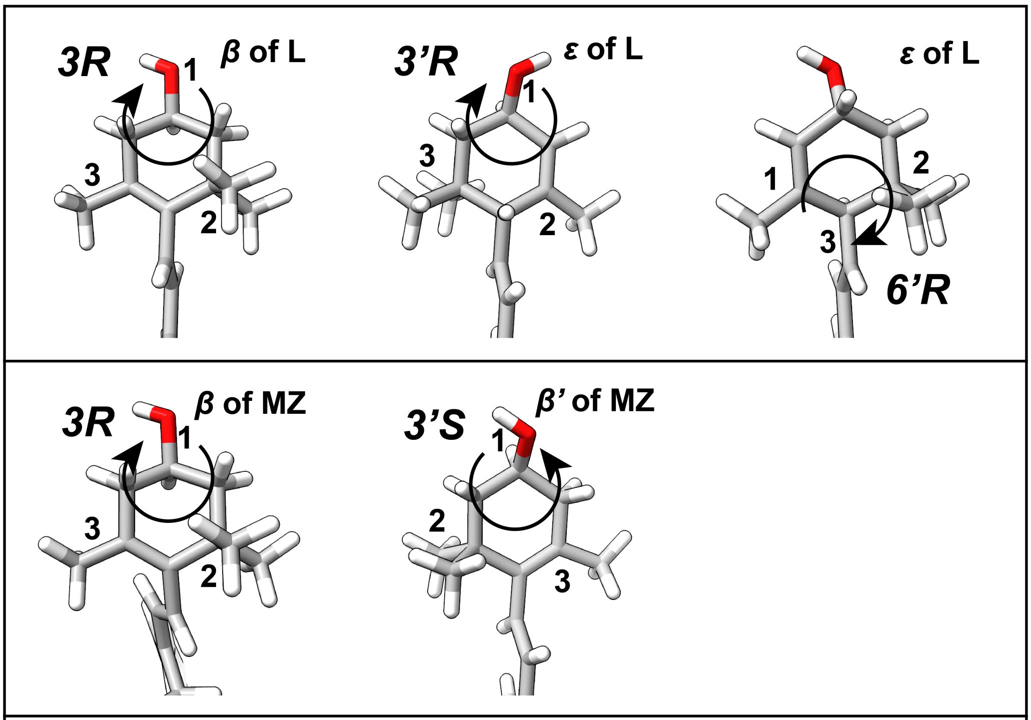


**Figure S3**. Priority assignment and stereoconfiguration for the chiral centers in *β* and *ε* rings of dietary L (top) and MZ (bottom). The structure of L is from PDB ID 6jo5 after correction of stereoconfiguration at C3. The structure of MZ was generated by merging ligands from PDB ID 7dz7 and PDB ID 1rwt. Note that C1, the carbon with two methyl groups has higher priority relative to C5, the carbon with one methyl group, for the *β* ring. But this priority ranking switches for the *ε* ring. Because of priority switching, C3 assigned as *R* in the *β* ring has opposite absolute handedness of C3' assigned as *R* in the *ε* ring. Moreover, upon π-bond migration, the C3' center of the resulting *β* ring in MZ is assigned *S* configuration despite having the same absolute handedness as C3' of the *ε* ring in L, which is assigned *R*.

Stereoconfiguration for LUT ligands found in the Protein Data Bank was corrected by mathematical operations to invert the C3 centers resulting in an ensemble of (*3R, 3'R, 6'R*) dietary L ligands. Coordinates for lutein ligands were obtained from structures of photosystems and light harvesting complexes in the Protein Data Bank. Specifically, all 122 LUT ligands from PDB IDs 1rwt, 4xk8, 5xnl, 6jo5, 6kac, and 7dz7 provided starting coordinates. Inspection of these starting coordinates showed 3S, 3'R, 6'R configuration for all LUT ligands. To convert these ligands to the correct *3R, 3'R, 6'R* configuration, the positions of atoms in the β ionone ring were reflected across the plane defined by the double bond connecting atoms C6 and C5. Atom reflection operations were executed by vector algebra, including cross product and dot product operations, as encoded in an R program *dietary.lutein.R* written by MPH. Briefly (x,y,z) coordinates for each atom in the β ionone ring (C1, C2, C3, O3, C4, C5, C6, C16, C17, C18) defined corresponding vectors. These vectors will be denoted here as the atom name in bold. For example the (x,y,z) vector defining the position of atom C1 is **C1**. The cross product (**C1** - **C6**) x (**C5** - **C6**) defined **R**, a vector perpendicular to the reflecting plane defined by atoms C1, C5 and C6 (Equation S1).

$$\mathbf{R} = (\mathbf{C1} - \mathbf{C6}) \times (\mathbf{C5} - \mathbf{C6}) \qquad \text{Eq S1}$$

For each atom being reflected, its original position defined a difference vector relative to an atom in the reflecting plane, either C1 for the two methyl groups C17 and C18, or C6 for all other atoms. The dot product of this difference vector with **R** provided the distance from the reflecting plane, and thereby a means to compute the projection vector along **R**. The coordinates of each atom could then be transformed by subtracting this projection vector scaled by 2. Equations below illustrate these computations for reflection of atom O3. Similar calculations were applied to all atoms in the β ionone ring, except those atoms that are already in the reflecting plane (C1, C4, C5, C18).

$$\text{distance}(\mathbf{O3}) = \mathbf{R} \bullet (\mathbf{O3} - \mathbf{C6}) \qquad \text{Eq. S2}$$

$$\text{projection}(\mathbf{O3}) = \mathbf{R}\ \text{distance}(\mathbf{O3}) / \mathbf{R} \bullet \mathbf{R} \qquad \text{Eq. S3}$$

$$\text{reflection}(\mathbf{O3}) = \mathbf{O3} - 2\ \text{projection}(\mathbf{O3}) \qquad \text{Eq. S4}$$

The resulting ensemble of 122 dietary L ligands was processed with the *prepare_ligand4.py* module of AutoDock Tools (ADT) with the options -A hydrogens -U nphs_lps -Z, as executed by a *tcsh* shell script *pdb2pdbqt-dietarylutein.script* written by author MPH. To create versions of these L ligands with rotatable bonds connecting ionone rings to the polyene chain, the *tcsh* script piped a set of *vi* commands to the *vi* editor that modified the text in a duplicate .pdbqt file. These commands are encoded in *activate_torsionC6C7C26C27_lut.vi*, a *vi* script also written by the author MPH.

**Creating the ensemble of MZ and Z ligands**

The coordinates for (*3R, 3'S*) MZ ligands and (*3R, 3'R*) Z ligands were constructed by joining portions of two xanthophylls. For creating the ensemble of MZ ligands, each member of the dietary L ensemble, the *ε* ring was replaced with a *β* ring randomly selected from the original LUT ensemble with *3S* configuration. (*3R, 3'R*) Z ligands were similarly constructed by replacing the *ε* ring found in the dietary L ensemble with a *β* ring randomly selected from the dietary L molecules with *3R* configuration. In these *ε* ring replacement operations, the second *β* ring was positioned as closely as possible with the *ε* ring by superposition of C7, C8 and C9 atoms of the polyene chain of the second L or LUT molecule onto C27, C28 and C29 atoms in the first L molecule. Coordinates for the hybrid ligand molecule were written by selectively merging C1, C2, C3, O3, C4, C5, C6, C7, C8, C9, C10, C11, C12, C13, C14, C15, C16, C17, C18, C19, C20, C29, C30, C31, C32, C33, C34, C35, C39, C40 atoms belonging to the *β* ring and polyene chain from the first L molecule, thereby excluding the atoms belonging to the *ε* ionone ring, with the C1, C2, C3, O3, C4, C5, C6, C7, C8, C16, C17, C18 atoms from the *β* ionone ring of the second L or LUT molecule after superposition. Superposition and the merging of atoms was facilitated by ChimeraX scripts, *superimpose.Sbeta.to.epsilon.cxc* for MZ ligands and by *superimpose.beta.to.epsilon.cxc* for Z ligands. Further cleanup of atom names was accomplished with a *tcsh* script *cleanup.mesozeaxanthin.script* and a set of *vi* commands encoded with *cleanup.mesozeaxanthin.vi* for the MZ ligands, and a *tcsh* script *cleanup.zeaxanthin.script* and a set of *vi* commands encoded with *cleanup.zeaxanthin.vi* for the Z ligands. These scripts for superposition and for atom name cleanup were created by an R program *dietary.zeaxanthin.R* written by the author MPH..

MZ and Z ligands were prepared for docking by processing with the *prepare_ligand4.py* module of AutoDock Tools (ADT) with the options -A hydrogens -U nphs_lps -Z, as executed by *tcsh* shell scripts *pdb2pdbqt-mesozeaxanthin.script* and *pdb2pdbqt-dietaryzeaxanthin.script* written by author MPH. To create versions of these MZ and Z ligands with rotatable bonds connecting ionone rings to the polyene chain, the *tcsh* script piped a set of *vi* commands to the *vi* editor that modified the text in duplicate pdbqt files. These commands are encoded in the scripts *activate_torsionC6C7C26C27_mesozea.vi* and *activate_torsionC6C7C26C27_zea.vi* written by the author MPH.

**Removal of identical ligands**

Ensembles for L, Z and MZ ligands were examined to identify identical ligands with R.M.S.D. less than ~0.01 Å after superposition. Redundant ligands were excluded, resulting in ensembles with 93 unique dietary L, 119 MZ, and 121 dietary M ligands.

**Ensemble of β-carotene ligands**

Coordinates for *β*-carotene ligands were derived from the BCR ligands found in structures of photosystems determined by x-ray crystallography or cryo-electron microscopy to a resolution limit of 2.2 Å or better, as archived with the Protein Data Bank. Specifically, all BCR ligands from PDB IDs 3wu2, 4ub6, 5b5e, 5mx2, 5v2c, 6jLj, 6w1o, and 7d1u provided the coordinates for 170 β-carotene ligands. This initial ensemble was examined to identify identical conformations, as judged by R.M.S.D less than ~0.01 Å after superposition. Redundant ligands were removed, resulting in an ensemble of 84 non-redundant *β*-carotene ligands. *β*-carotene ligands were prepared for docking by processing with the *prepare_ligand4.py* module of AutoDock Tools (ADT) with the options -A hydrogens -U nphs_lps -Z, as executed by a *tcsh* shell scripts *pdb2pdbqt-betacarotene.script* written by author MPH.

**Molecular docking**

We applied *Autodock VINA* [29,30] to dock members of the dietary L ensemble as rigid ligands to bovine RPE65 protein (PDB ID 7L0e) [36]. To alleviate steric clashes, three protein residues lining the substrate binding tunnel were allowed to adopt altered rotamers. These flexible residues were Phe61, Val134 and Tyr338. The flexible residues were separated from the rigid receptor structure with use of *Autodock Tools* [63,64]. The search volume was constrained to the portion of the RPE65 structure where competitive inhibitors for RPE65 are found in crystal structures, specifically defined by the parameters: center_x= -40.6, center_y= 138.6, center_z= 11.0, size_x= 24.0, size_y= 25.5, size_z= 17.25. Docking was performed with gpu or cpu nodes available through the Center for High Performance Computing at the University of Utah.

Batch jobs for docking with VINA were automated with the SLURM scripts *vina_STQ2.slurm* for L, *vina_VFQ2.slurm* for MZ, *vina_8UH2.slurm* for Z, and *vina_BCR2.slurm* for *β*-carotene. The top 5 poses obtained for each ligand were isolated with the *vina_split* module of AutoDock VINA and energy terms were obtained by running AutoDock VINA with the *-score_only* option. Ligand isolation and energy term extraction was performed with gpu or cpu nodes available through the Center for High Performance Computing at the University of Utah. Batch jobs were automated with the SLURM scripts: *split_scoreonly_RPE65_STQ2.slurm* for L, *split_scoreonly_RPE65_VFQ2.slurm* for MZ, *split_scoreonly_RPE65_8UH2.slurm* for Z, and *split_scoreonly_RPE65_BCR2.slurm* for *β*-carotene. The docking outcomes and binding energies were analyzed with *Binding.Energy.Analysis.RPE65-2025.Rsource.R*, an R program written by MPH. Molecular docking outcomes were displayed with ChimeraX [65].
